## Supplementary Information for "Light inducible protein degradation in *E. coli* with the LOVdeg tag"

#### Supplementary Tables

**Table S1**. Plasmids used in this study.

| ***Plasmid*** | ***Origin*** | ***Operon*** | ***Resistance*** | ***Reference*** |
|---|---|---|---|---|
| *pBbS5c-mCherry-AsLOV2(546)* | SC101 | $P_{lacUV5}$-mCherry-AsLOV2(546) | $Cm^R$ | This study |
| *pBbS5c-mCherry-AsLOV2(543)* | SC101 | $P_{lacUV5}$-mCherry-AsLOV2(543) | $Cm^R$ | This study |
| *pBbS5c-mCherry-AsLOV2*(546)* | SC101 | $P_{lacUV5}$-mCherry-AsLOV2*(546) | $Cm^R$ | This study |
| *pBbS5c-mCherry-AsLOV2*(543)* ***(LOVdeg)*** | SC101 | $P_{lacUV5}$-mCherry-AsLOV2*(543) | $Cm^R$ | This study |
| *pBbSW7c-mCherry-AsLOV2*(543)* | SC101 | $P_{const.}$-mCherry-AsLOV2*(543) | $Cm^R$ | This study |
| *pBbA8k-ClpA* | P15a | $P_{Bad}$-ClpA | $Kan^R$ | This study |
| *pBbA8k-HslUV* | | | | |
| | P15a | $P_{bad}$-HslVU | $Kan^R$ | This study |
| *pBbS5c-LacI-LOVdeg-mCherry* | SC101 | $P_{lac}$-LacI-LOVdeg, $P_{lacUV5}$-mCherry | $Cm^R$ | This study |
| *pBbAa-LacI-decoy* | P15a | LacI decoy | $Amp^R$ | (Wang et al., 2021) |
| *pJF076Sk* | SC101 | $P_{J23117}$-mRFP1 | $Kan^R$ | This study |
| *pJF093* | P15a | $P_{Cas9}$-dCas9, $P_{tet}$-MCP-SoxS | $Cm^R$ | (Dong et al., 2018) |
| *pA1c-CRISPRa* | P15a | $P_{Cas9}$-dCas9, $P_{trc}$-MCP-SoxS | $Cm^R$ | This study |
| *pA1c-CRISPRa-LOVdeg* | P15a | $P_{Cas9}$-dCas9, $P_{trc}$-MCP-SoxS-LOVdeg | $Cm^R$ | This study |
| *pCD061-J106* | ColE1 | $P_{J23119}$-J106-scRNA | $Amp^R$ | (Dong et al., 2018) |
| *pBbA5k-acrAB* | P15a | $P_{lacUV5}$-acrAB | $Kan^R$ | (El Meouche and Dunlop, 2018) |
| *pBbA5k-acrAB-LOVdeg* | P15a | $P_{lacUV5}$-acrAB-LOVdeg | $Kan^R$ | This study |
| *pBbS5c-mCherry-LOVdeg(V416I)* | SC101 | $P_{lacUV5}$-mCherry-LOVdeg(V416I) | $Cm^R$ | This study |
| *pEL222* | P15a | $P_{rrnBp1}$-EL222 | $Cm^R$ | (Jayaraman et al., 2016) |
| *pBbE5k-$P_{EL222}$-mCherry-LOVdeg* | ColE1 | $P_{EL222}$-mCherry-LOVdeg | $Kan^R$ | This study, promoter from (Ding et al., 2020) |
| *pBbS5c-CpFatB1*-LOVdeg* | SC101 | $P_{lacUV5}$-*Cp*FatB1.2-M4-287-LOVdeg | $Cm^R$ | This study |
| *pBbE5k-$P_{EL222}$-CpFatB1*-LOVdeg* | ColE1 | $P_{EL222}$- *Cp*FatB1.2-M4-287-LOVdeg | $Kan^R$ | This study |

**Table S2**. Strains used in this study.

| *Strain* | *Relevant genotype* | *Reference* |
|---|---|---|
| *BW25113 (Wild type)* | F$^-$ Δ(araD-araB)567 ΔlacZ4787(::rrnB-3) λ$^-$ rph-1 Δ(rhaD-rhaB)568 hsdR514 | (Baba et al., 2006) |
| *ΔlacI* | *E. coli* BW25113 Δ*lacI*, cured from Keio collection | (Baba et al., 2006) |
| *ΔacrB* | *E. coli* BW25113 Δ*acrB*, cured from Keio collection | (Baba et al., 2006) |
| *ΔclpX* | *E. coli* BW25113 Δ*clpX*, cured from Keio collection | (Baba et al., 2006) |
| *ΔhslU* | *E. coli* BW25113 Δ*HslU*, cured from Keio collection | (Baba et al., 2006) |
| *ΔclpA* | *E. coli* BW25113 Δ*clpA,* cured from Keio collection | (Baba et al., 2006) |
| *Δlon* | *E. coli* BW25113 Δ*lon*, cured from Keio collection | (Baba et al., 2006) |
| *ΔclpS* | *E. coli* BW25113 Δ*clpS*, cured from Keio collection | (Baba et al., 2006) |
| *ΔclpP* | *E. coli* BW25113 Δ*clpP*, cured from Keio collection | (Baba et al., 2006) |
| *ΔacrB + AcrAB* | Δ*acrB*, pBbA5k-acrAB | (El Meouche and Dunlop, 2018) |
| *ΔacrB + AcrAB-LOVdeg* | Δ*acrB*, pBbA5k-acrAB-LOVdeg | This study |
| *LacI-LOVdeg* | Δ*lacI*, pBbS5c-LacI-LOVdeg-mCherry | This study |
| *LacI-LOVdeg + decoy* | Δ*lacI*, pBbS5c-LacI-LOVdeg-mCherry | This study |
| *Dong et al. CRISPRa* | BW25113, pJF093, pJF076Sk, pCD061-J106 | (Dong et al., 2018) |
| *$P_{trc}$-inducible CRISPRa* | BW25113, pA1c-CRISPRa, pJF076Sk, pCD061-J106 | This study |
| *LOVdeg CRISPRa* | BW25113, pA1c-CRISPRa-LOVdeg, pJF076Sk, pCD061-J106 | This study |
| *LOVdeg only ($P_{EL222}$)* | BW25113, pBbE5k-$P_{EL222}$-mCherry-LOVdeg | This study |
| *LOVdeg + EL222* | BW25113, pBbE5k-$P_{EL222}$-mCherry-LOVdeg, pEL222 | This study |
| *CpFatB1*-LOVdeg* | BW25113, pBbS5c-CpFatB1*-LOVdeg | This study |
| *CpFatB1*-LOVdeg + EL222* | BW25113, pBbE5k-$P_{EL222}$-CpFatB1*-LOVdeg, pEL222 | This study |
| *CpFatB1* EL222* | BW25113, pBbE5k-$P_{EL222}$-CpFatB1*-LOVdeg | This study |

**Table S3**. DNA sequences used in this study.

| ***Gene*** | ***Sequence*** |
|---|---|
| *AsLOV2(546)* | ttggctactacacttgaacgtattgagaagaactttgtcattactgacccaagattgccagataatcccattatattcgcgtccgatagt<br>ttcttgcagttgacagaatatagccgtgaagaaattttgggaagaaacTGCaggtttctacaaggtcctgaaactgatcgcgcga<br>cagtgagaaaaattagagatgccatagataaccaaacagaggtcactgttcagctgattaattatacaaagagtggtaaaaagttct<br>ggaacctctttcacttgcagcctatgcgagatcagaagggagatgtccagtactttattggggttcagttggatggaactgagcatg<br>tccgagatgctgccgagagagagggagtcatgctgattaagaaaactgcagaaaatattgatgaggcggcaaaagaactt |
| *AsLOV2(543)* | ttggctactacacttgaacgtattgagaagaactttgtcattactgacccaagattgccagataatcccattatattcgcgtccgatagt<br>ttcttgcagttgacagaatatagccgtgaagaaattttgggaagaaacTGCaggtttctacaaggtcctgaaactgatcgcgcga<br>cagtgagaaaaattagagatgccatagataaccaaacagaggtcactgttcagctgattaattatacaaagagtggtaaaaagttct<br>ggaacctctttcacttgcagcctatgcgagatcagaagggagatgtccagtactttattggggttcagttggatggaactgagcatg<br>tccgagatgctgccgagagagagggagtcatgctgattaagaaaactgcagaaaatattgatgaggcggca |
| *AsLOV2*(546)* | ttggctactacacttgaacgtattgagaagaactttgtcattactgacccaagattgccagataatcccattatattcgcgtccgatagt<br>ttcttgcagttgacagaatatagccgtgaagaaattttgggaagaaactgtcgcttcttacaaggcccggaaaccgatcgcgcaac<br>cgtccgtaagattcgcgacgccatcgataatcaaaccgaggttacggtgcaattaattaactacacgaaatccggtaagaagttttg<br>gaatgtatttcatttgcaacccatgcgtgaccagaaaggagatgtacaatatttcatcggagttcaactcgacggtacggagcgcct<br>tcgcggggcagcggaacgcgaagccgttatgttgattaagaaaaccgcgttccaaatcgcggaggcggcaaaagaactt |
| *AsLOV2*(543)*<br>**LOVdeg** | ttggctactacacttgaacgtattgagaagaactttgtcattactgacccaagattgccagataatcccattatattcgcgtccgatagt<br>ttcttgcagttgacagaatatagccgtgaagaaattttgggaagaaactgtcgcttcttacaaggcccggaaaccgatcgcgcaac<br>cgtccgtaagattcgcgacgccatcgataatcaaaccgaggttacggtgcaattaattaactacacgaaatccggtaagaagttttg<br>gaatgtatttcatttgcaacccatgcgtgaccagaaaggagatgtacaatatttcatcggagttcaactcgacggtacggagcgcct<br>tcgcggggcagcggaacgcgaagccgttatgttgattaagaaaaccgcgttccaaatcgcggaggcggca |

### Supplementary Figures

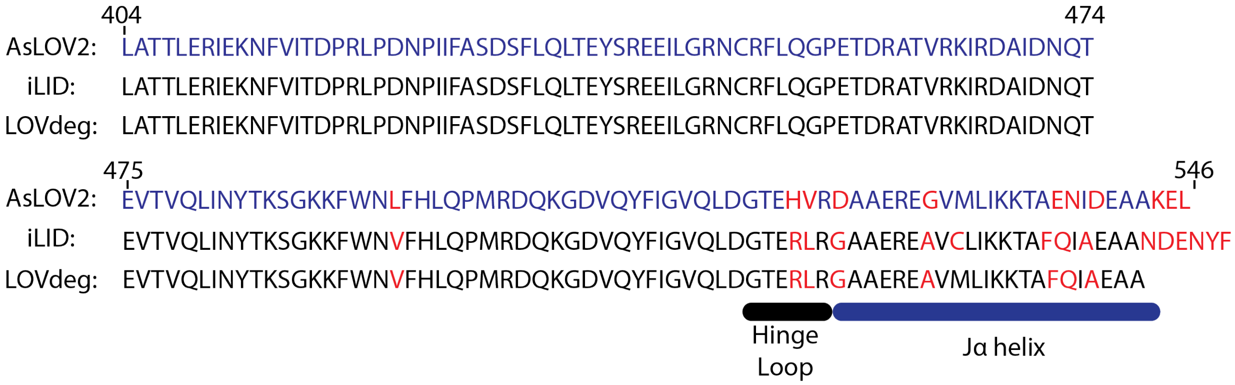


**Figure S1**. Alignment of *As*LOV2, iLID (mutated version of *As*LOV2 from Guntas *et al*. (Guntas et al., 2015)) and LOVdeg (*As*LOV2*(543)) amino acid sequences.

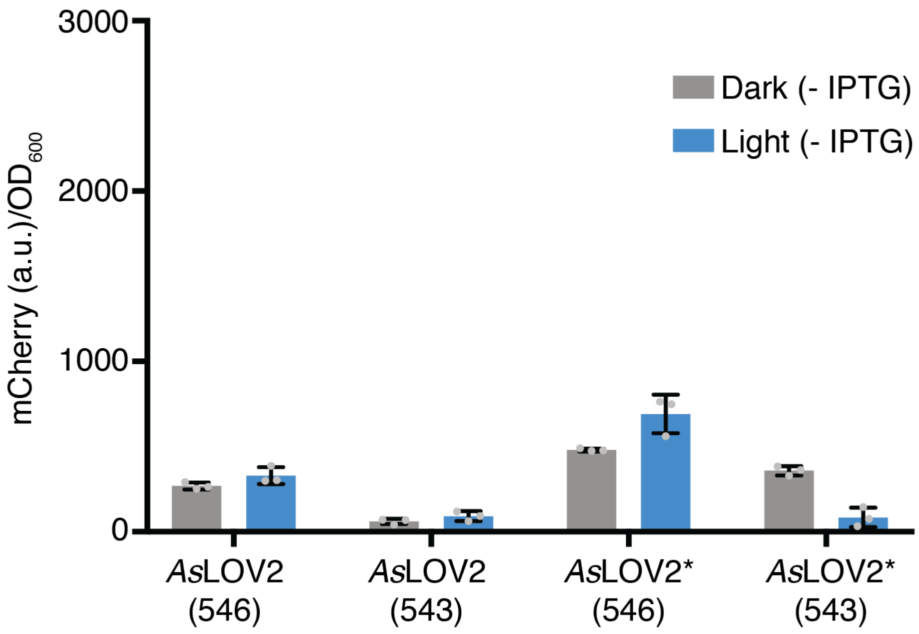


**Figure S2**. mCherry expression levels without IPTG induction in response to 465 nm blue light for wild type *As*LOV2 and mutated *As*LOV2* fusions with and without truncation. Error bars show standard deviation around the mean (n=3 biological replicates).

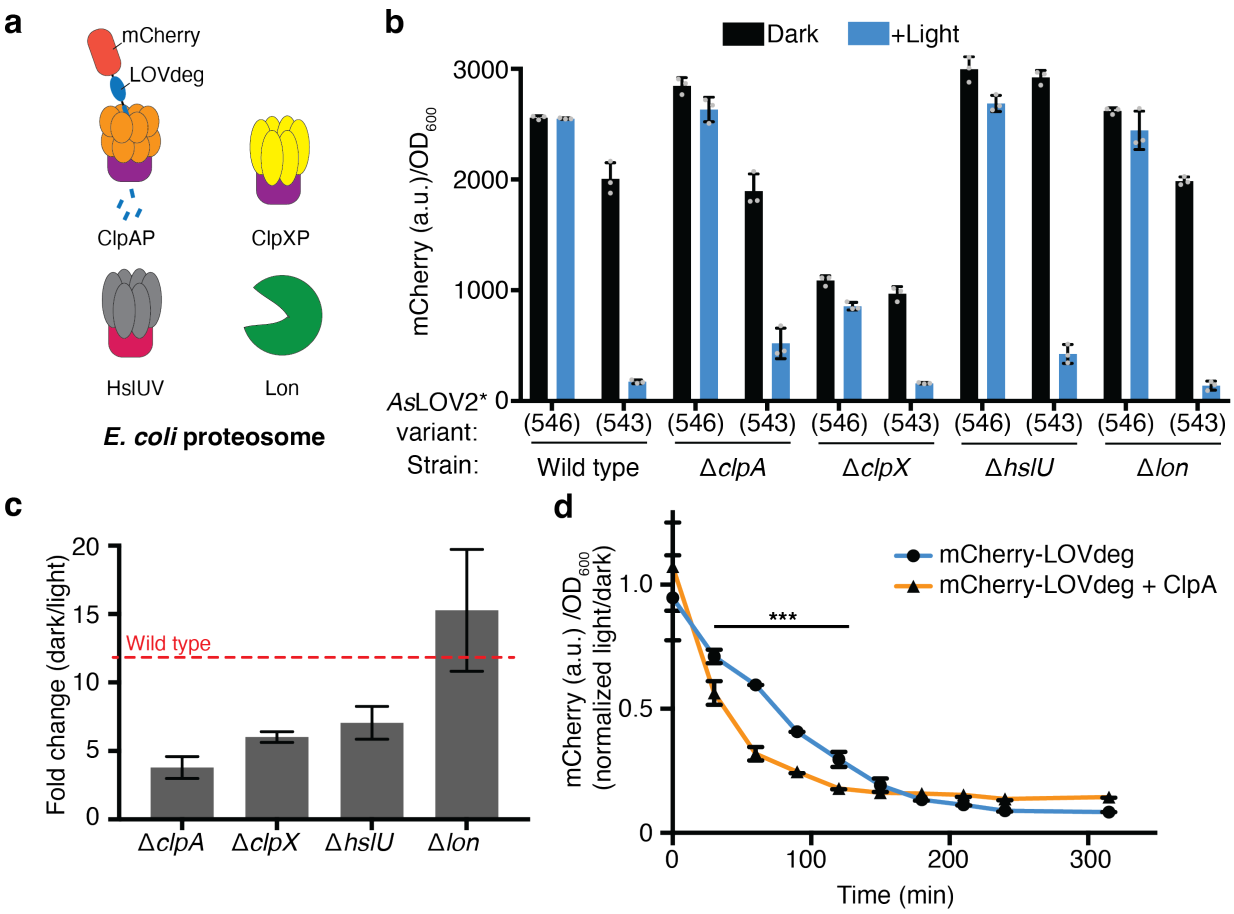


**Figure S3.** Investigating proteasome components involved in LOVdeg tag destabilization. **(a)** Unfoldases and proteases of the *E. coli* proteasome. **(b)** Light-dependent stability of constitutively expressed mCherry fusions with truncated (*As*LOV2*(543), LOVdeg) and non-truncated (*As*LOV2*(546)) tags in strains lacking endogenous unfoldases or proteases. **(c)** Fold change degradation of mCherry-LOVdeg (i.e. mCherry- *As*LOV2*(543)) in strains lacking endogenous unfoldases. Fold change compares ratio of dark to light states. **(d)** Expression of mCherry-LOVdeg over time under light exposure in wild type cells or cells overexpressing ClpA. Fluorescence signal is normalized to expression of cells kept in the dark (***$p < 0.0001$, comparison between data at the same time point, two tailed unpaired t-test). Error bars show standard deviation around the mean (n=3 biological replicates).

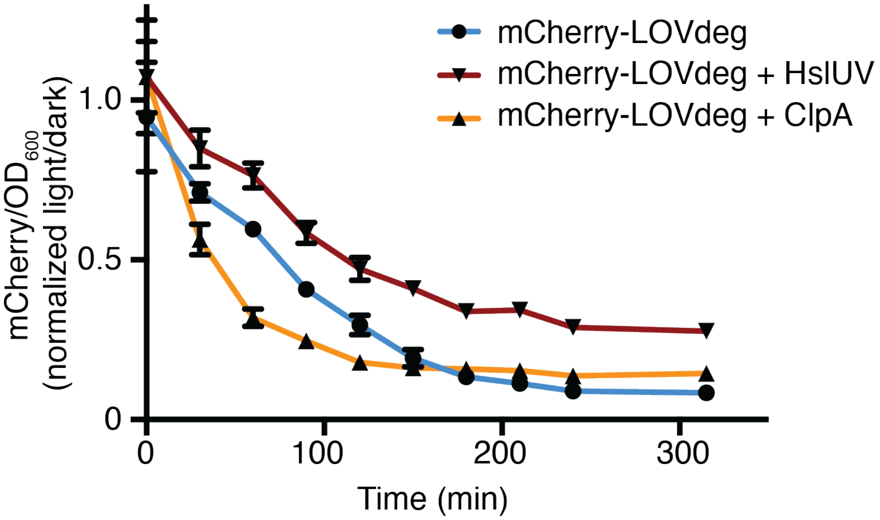


**Figure S4.** Expression of mCherry-LOVdeg over time under light exposure in wild type cells, cells expressing exogenous HslUV, and cells expressing exogenous ClpA. Fluorescence signal is normalized to expression of cells kept in the dark. Error bars show standard deviation around the mean (n=3 biological replicates). ClpA expression data is the same as that shown in Fig. S3 and included for comparison.

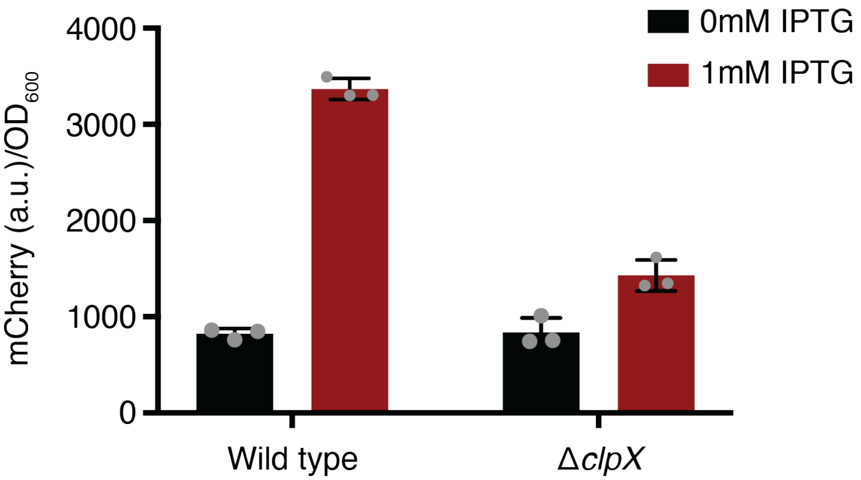


**Figure S5**. Untagged mCherry expression induced with IPTG in wild type and *clpX* knockout strains. Error bars show standard deviation around the mean (n=3 biological replicates).

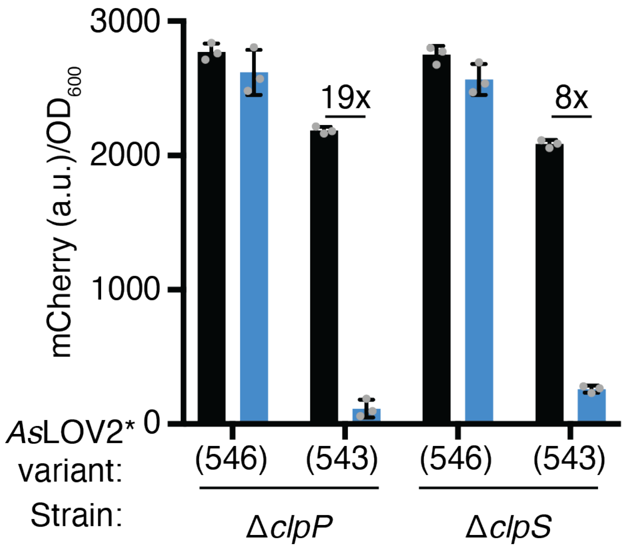


**Figure S6.** Light-dependent stability of mCherry fusions with truncated and non-truncated LOVdeg tags in strains lacking *clpP* and *clpS*. Error bars show standard deviation around the mean (n=3 biological replicates).

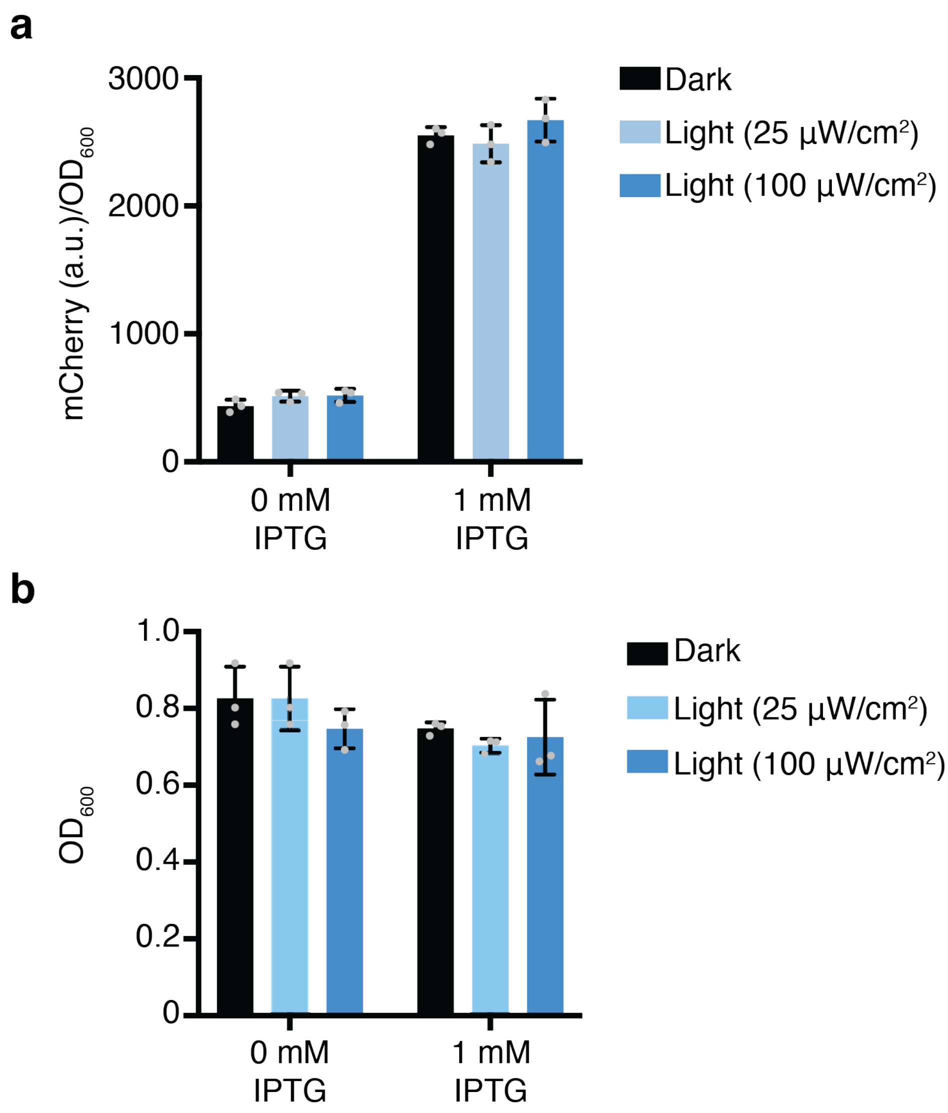


**Figure S7.** Light response of mCherry without any *As*LOV2 variant. **(a)** mCherry protein levels in response to dark or 465 nm blue light. (b) Culture densities ($OD_{600}$) of strains from (a). Error bars show standard deviation around the mean (n = 3 biological replicates).

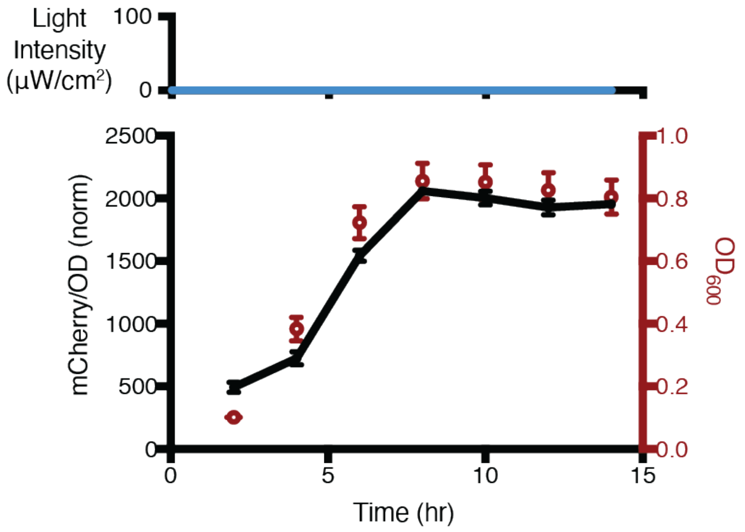


**Figure S8.** mCherry-LOVdeg fluorescence levels and growth without any light exposure. This curve is used to normalize data in Fig. 1f at each time point so that it is possible to see relative decreases in protein levels. Error bars show standard deviation around the mean (n = 3 biological replicates).

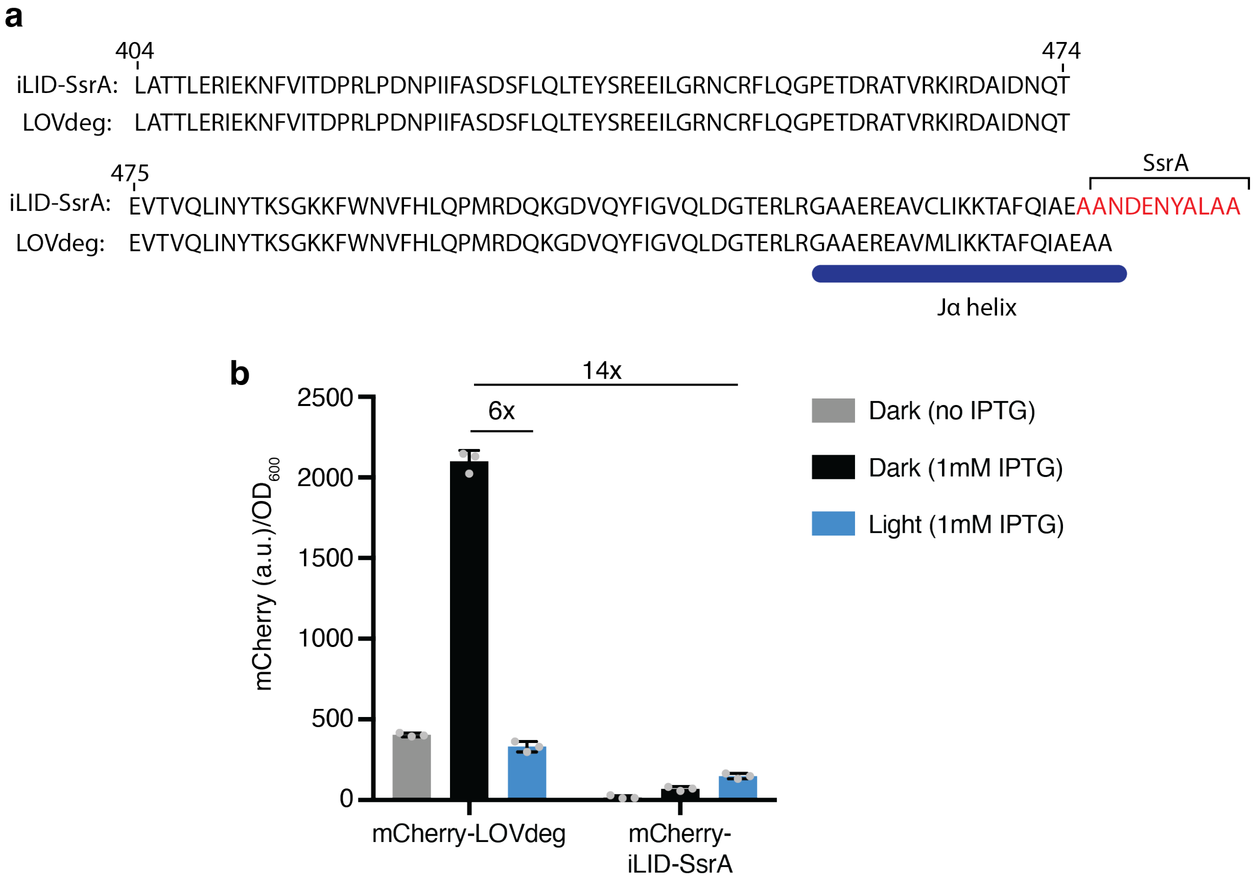


**Figure S9. (a)** Alignment of iLID modified to contain a full length SsrA tag and the LOVdeg tag. **(b)** Protein level comparison between mCherry-LOVdeg and an analog with a constitutively active SsrA tag. Error bars show standard deviation around the mean (n = 3 biological replicates).

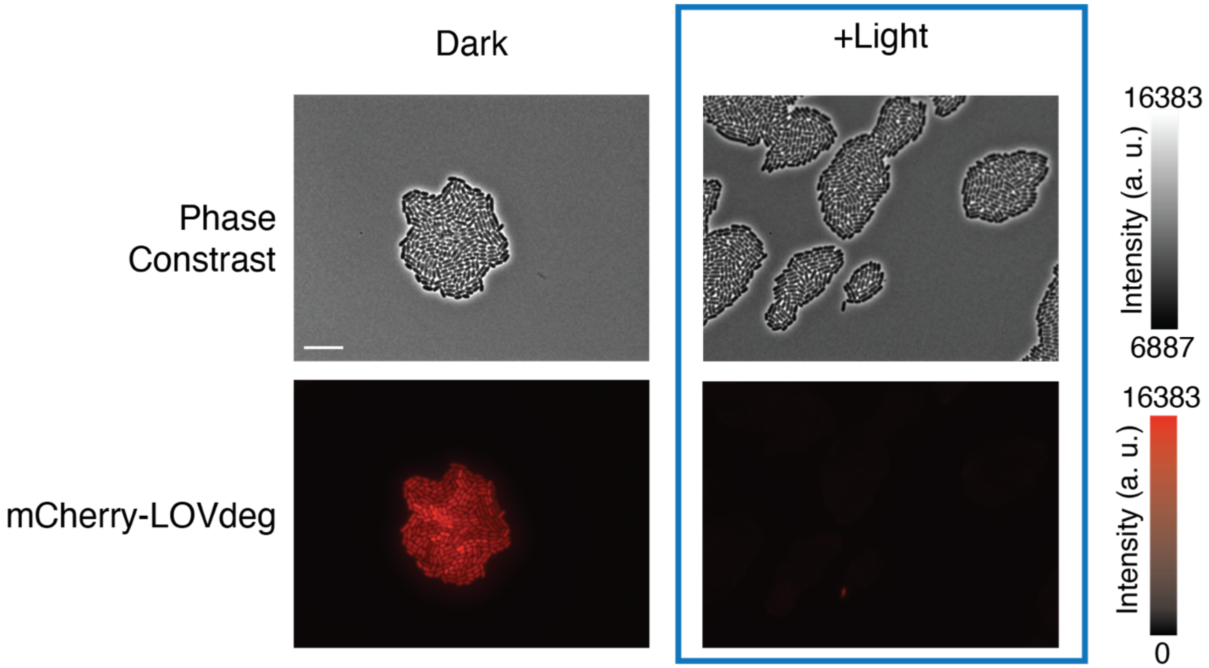


**Figure S10.** Phase contrast and fluorescence images of cells constitutively expressing mCherry-LOVdeg exposed to blue light or kept in the dark (Scale bar = 10 μm).

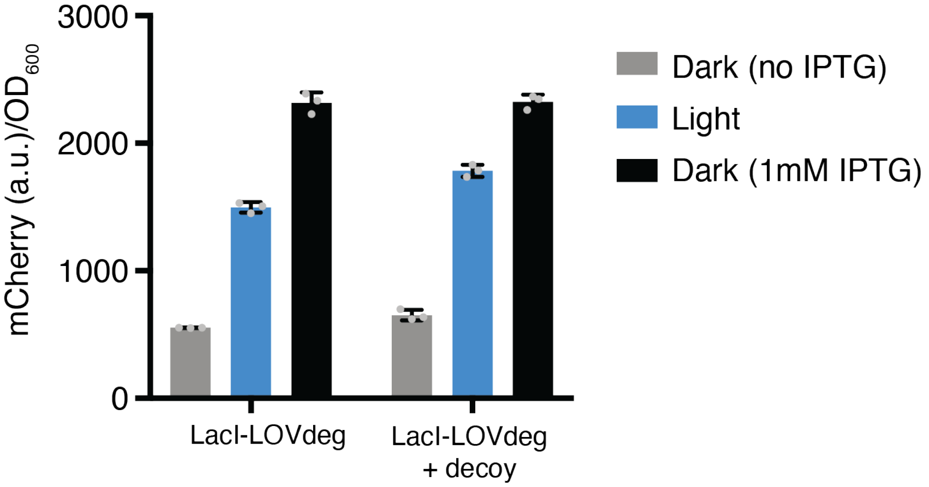


**Figure S11**. LacI-LOVdeg and LacI-LOVdeg + decoy control of mCherry with 1 mM IPTG induction included. Error bars show standard deviation around the mean (n=3 biological replicates).

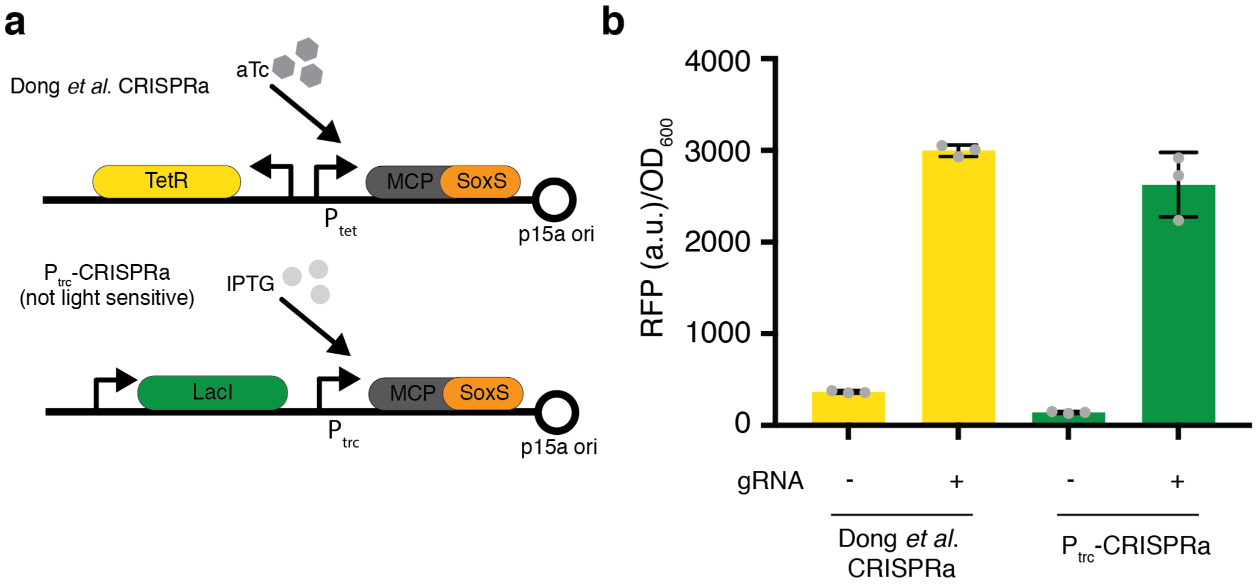


**Figure S12. (a)** Genetic constructs of the original SoxS-CRISPRa from Dong *et al*. where the activator is aTc inducible. However, aTc is light sensitive, thus we replaced the TetR/$P_{tet}$ portion with the IPTG-inducible LacI/$P_{trc}$. **(b)** Comparison of the CRISPRa activity of the original construct and the new $P_{trc}$ promoter construct with and without a gRNA targeting the promoter of mRFP1. Error bars show standard deviation around the mean (n=3 biological replicates).

### Supplementary Text

*Mechanistic insights into the E. coli proteasome*

Based on the exposed C-terminal amino acids in the LOVdeg (E-A-A), we anticipated that degradation would be primarily mediated by ClpXP because the C-terminal alanines, when on an unstructured peptide of sufficient length, are known to be adequate for ClpX targeting (Fei et al., 2020). However, degenerate degradation from multiple endogenous proteases is common in *E. coli*, and has been demonstrated with SsrA tagged substrates (Flynn et al., 2001). Additionally, ClpX is the most extensively studied unfoldase, meaning the rules governing other endogenous unfoldases are less understood and their action should not be ruled out. ClpA, ClpX, and HslU unfoldases utilize protease counterparts to perform protein degradation with ClpAP, ClpXP, and HslVU complexes, respectively. In contrast, Lon performs both unfoldase and protease activity. To understand the endogenous unfoldase(s) responsible for degradation of the LOVdeg tag, we expressed the mCherry-LOVdeg construct in several unfoldase knockouts, including those deleting the genes that encode ClpA, ClpX, HslU, or Lon (Fig. S3a). FtsH, the last of the five unfoldase-proteases in *E. coli*, was not included in the knockout study because it is an essential gene and could not be knocked out.

We measured reductions in mCherry in response to blue light induction in the different knockout backgrounds. In each knockout, we expressed mCherry-LOVdeg under a constitutive promoter. We also expressed the non-truncated counterpart for a degradation-resistant comparison. Protein expression was decreased in light for all knockout strains, however the degree of reduction varied (Fig. S3b). The fold change of degradation was reduced relative to wild type in Δ*clpA*, Δ*clpX*, and Δ*hslU* strains (Fig. S3c). To further test which of these unfoldases was responsible for LOVdeg tag degradation, we created plasmids expressing each unfoldase exogenously under an arabinose inducible promoter to see if excess unfoldase would increase degradation of mCherry. ClpA was the only one to display increased degradation when overexpressed (Fig. S3d). With ClpA expressed from plasmid in addition to endogenous ClpA, the half-life of mCherry-LOVdeg was decreased from 74 to 38 minutes. Given the fold change decrease we observed in Δ*hslU*, we were surprised that strains overexpressing HslU did not increase their degradation rate (Fig. S4). The lack of enhanced degradation when HslUV is overexpressed suggests that it is not the primary source of LOVdeg tag degradation. It is possible that the decreased degradation fold change seen in Δ*hslU* can be attributed to broader systemic changes in this knockout strain. Although the Δ*clpX* strain showed a reduced fold change, this is likely due to generalized changes in expression, which we observed with mCherry with no degradation tag in this strain as well (Fig. S5). Our initial assumption that ClpXP would be the primary source of degradation was incorrect. The data instead show that ClpA is involved in LOVdeg tag degradation, however, it is likely a single unfoldase-protease is not entirely responsible for degradation.

Since ClpA was implicated in degradation, we also tested knockouts of ClpP and ClpS. ClpP is responsible for proteolysis of substrates unfolded by ClpA, and ClpS is an adaptor protein for ClpAP that alters targeting specificity (Baker and Sauer, 2006). mCherry was degraded efficiently in both the Δ*clpP* and Δ*clpS* strains (Fig. S6). Because ClpA is implicated in degradation, we initially expected ClpP, the protease counterpart to ClpA, would be necessary. However, studies examining degradation of SsrA tagged substrates have shown that substrates can still be degraded efficiently, even in Δ*clpP* strains (Farrell et al., 2005; Lies and Maurizi, 2008). For example, Lies *et al*. found that SsrA tagged substrates can be degraded in Δ*clpP* strains but accumulate in Δ*clpP* Δ*lon* strains. They concluded that in the absence of ClpP, ClpA and ClpX continue to unfold substrates and Lon carries out proteolysis on the unfolded substrates. A similar mechanism may be at play with the LOVdeg tag in Δ*clpP* cells.

By knocking out exogenous *E. coli* unfoldases, we gained partial insight into the mechanism of LOVdeg tag destabilization. The Δ*clpA* knockout exhibits decreased degradation in response to light, while complementing cells with *clpA* increases degradation speed, demonstrating the involvement of the ClpA protease in LOVdeg tag destabilization. However, other proteolytic activity is also involved, as degradation could still be achieved, albeit to a lesser extent, in the Δ*clpA* strain. Further, full degradation was maintained in its partner Δ*clpP* strain. It remains unclear whether the LOVdeg tag is targeted due to specific amino acid sequence interactions with a given unfoldase or if the general disorder induced at the C-terminal end of the protein is sufficient for recognition by the proteasome. The non-truncated versions of *As*LOV2 and *As*LOV2*, which maintain light dependent C-terminal disorder, are stable, suggesting that it is a mix of sequence and C-terminal peptide disorder. The full mechanism of LOVdeg tag destabilization is a topic for future investigation.

*Adding decoy sites to reduce impact of basal LacI does not improve LacI-LOVdeg response*

We sought to further investigate why LacI-LOVdeg responded with higher mCherry expression under IPTG induction than light induction. One possibility is that low levels of LacI are escaping degradation and causing basal repression. To address this, we added LacI decoy binding sites that work by binding excess LacI, following a method we developed in a previous study (Wang et al., 2021). With the decoy present, light induced expression was only slightly increased (Fig. S11). This indicates that low levels of LacI are not the primary explanation for the discrepancy between IPTG induction and light induction. We hypothesize that the lowered expression with light exposure stems from the time delay inherent in protein degradation compared to allosteric binding of IPTG to LacI.
